# Genetic Mapping of Progressive Ethanol Consumption in the Diversity Outbred Mouse

**DOI:** 10.1101/2022.12.06.519344

**Authors:** Zachary Tatom, Kristin M. Mignogna, Zachary Sergi, Jeremy Nguyen, Marie Michenkova, Maren L. Smith, Michael F. Miles

## Abstract

Traditional genetic mapping studies using inbred crosses are a powerful tool for identifying chromosomal regions associated with ethanol-related traits, but typically have very large confidence intervals which make identification of specific and potentially causal candidate genes difficult. Diversity Outbred (DO) mice offer the ability to map quantitative trait loci (QTLs) associated with ethanol-drinking behaviors at a high resolution that allows for easier identification of candidate genes. Here, we exposed a population of 636 male DO mice to four weeks of intermittent ethanol access via a three-bottle choice paradigm, identifying 3 significant (Chrs 3, 4, and 12) and 12 suggestive loci for ethanol-drinking behaviors. The confidence intervals for these loci were narrow (1-4 Mbp for significant QTLs). We then further analyzed positional candidate genes using transcriptomics data from prefrontal cortex samples taken from 220 of these animals, as well as human GWAS data and prior gene set data for ethanol or other drugs of abuse. These results represent the highest-resolution genetic mapping of ethanol consumption behaviors in mice to date, providing for the identification of novel loci and candidate genes for progressive ethanol consumption, including *Car8* --the lone gene with a significant *cis-*eQTL in strong linkage disequilibrium with our QTL for last week ethanol consumption on Chr 4.

## Introduction

Alcohol use disorder (AUD) poses a significant global healthcare burden, contributing to around 3 million deaths each year or roughly 5.3% of all deaths worldwide [1]. In the United States, 88,000 people die from causes related to alcohol use each year making it the third-largest cause of preventable death [2]. According to the National Survey on Drug Use and Health, 10.2% of people aged 12 or older in the United States met criteria for AUD in 2020; additionally, an estimated 22.2% of people reported binge alcohol use and 6.4% reported heavy alcohol use in 2020 [3]. Genetic factors are thought to play a significant role in alcohol consumption, with twin studies routinely estimating about 49% heritability in the risk for AUD [4].

Recent very large human genome-wide association studies (GWAS) have been employed to identify a limited number of genes and genetic loci associated with alcohol consumption [5, 6], dependence [7, 8], and AUD [9]. Significant environmental contributions to alcohol consumption and large sample sizes required to reach adequate power contribute to the difficulty in identifying genes from human GWAS. Animal models provide improved control over environmental variance and allow for eventual mechanistic and candidate gene identification studies infeasible in humans. To that end, mouse models have proven useful in identifying broadly-applicable biological mechanisms underlying specific behaviors associated with AUD including ethanol consumption [10], preference [11, 12], and withdrawal [13]. In particular, intermittent ethanol access (IEA) paradigms have been shown to model escalation of alcohol consumption as seen in the early stages of AUD in humans by consistently producing an increase in ethanol intake over time in mice [14]. Importantly, the IEA paradigm has been shown to vary across different mouse lines, suggesting that genetic variance impacting progressive consumption can be studied in such models [15].

Traditional ethanol behavioral quantitative trait loci (QTL) mapping in rodents often utilize short term F2 intercrosses, recombinant inbred lines produced by intercrosses between 2 inbred progenitor mouse strains, or similar strategies which lack genetic diversity and number of recombination events, typically leading to either poor statistical power to detect a QTL or broad QTL confidence intervals (>10-40 Mbp) with large numbers of potential candidate genes [16-18]. For example, a recent comprehensive study of ethanol binge drinking in BXD recombinant inbred mouse strains identified a number of suggestive loci but none reached statistical significance [16]. Therefore the task of using QTL studies to identify a tractable number of positional candidate genes that could be expeditiously verified as candidates modulating ethanol consumption was difficult until the advent of newer genetic resources [19].

Diversity Outbred (DO) mice were developed by Jackson Laboratories for high-resolution genetic mapping by utilizing a random outbreeding strategy to maintain high levels of allelic variation, recombination events, and heterogeneity [20]. This model was derived from 8 founder strains exhibiting a high degree of genetic and phenotypic diversity [21], capturing approximately 90% of the total genetic diversity of the mouse genome [22] and exhibiting diversity in ethanol-drinking phenotypes [23]. Recently, this model has been used to identify genetic loci corresponding to ethanol sensitivity, resulting in several novel loci with relatively narrow confidence intervals (∼4 Mbp) [24]. However, this novel mouse model has not yet been used to map QTLs for ethanol consumption behaviors.

Here we describe the first use of DO mice and an IEA paradigm to map behavioral QTLs (bQTLs) associated with progressive ethanol consumption. Our results show significant bQTLs on multiple chromosomes with generally narrow support intervals (most ranging from 1-5 Mbp). Positional candidate genes were then prioritized by a multi-step strategy using haplotype analysis, integration of DO mouse transcriptome data, and bioinformatics studies including merging of genetic data on AUD. Our results identify novel potential candidate genes and biological mechanisms for modulation of ethanol consumption behaviors. The work may both validate and broaden existing human studies on the genetics of alcohol consumption and AUD.

## Methods

### Ethics Statement

All animal care and euthanasia procedures were performed in accordance with the rules and regulations established by the United States Department of Agriculture Animal Welfare Act and Regulations, Public Health Services Policy on Humane Care and Use of Laboratory Animals, and American Association for Accreditation of Laboratory Animal Care. Humane endpoints were established by the same standards.

### Animal Housing and Power

Male DO mice (*n* = 636) were acquired from Jackson Laboratories after weaning at 4-6 weeks of age in 7 cohorts (average *n* = 106) spanning DO generations 22-25. Mice were singly-housed in temperature- and humidity-controlled vivariums on cedar shaving bedding with *ad libitum* access to water and standard chow (#7912, Harlan Teklad, Madison, WI, United States). Vivariums were set up with alternating 12-hour light and dark phases, and all mice were weighed weekly. Experimental procedures were begun on animals at 8-12 weeks of age.

The number of animals studied in this work was planned based on prior reports that estimated ∼600 DO mice were required to achieve 80% power to detect alleles causing less than 5% variance in a trait [25]. Female mice were excluded in this experiment because of likely sex-driven differences in ethanol consumption, including known differences in male and female ethanol consumption patterns in one of the DO progenitor strains, C57BL/6J. Including both sexes in this study would have required an untenable number of animals (n>1200) to achieve adequate genetic power.

### Intermittent Ethanol Access

Mice (*n* = 587) were exposed to ethanol via a three-bottle choice IEA paradigm for 5 weeks. (Figure 1) This consisted of alternating 24-hour periods of ethanol access (Monday, Wednesday, Friday), during which time mice were presented with three bottles (15% ethanol v/v, 30% ethanol v/v, and water). Ethanol was presented at the beginning of the dark phase and consumption was measured in mL 24 h later at the end of the following light phase. Bottles of each fluid were also placed on empty cages to correct for evaporation and spillage, and placement order of bottles was randomized each IEA day to avoid place preference confounds. Ethanol and water volume measurements were used to derive values for ethanol consumption (g ethanol/kg body weight per 24 hours), preference for ethanol compared to water (“ethanol preference”) and for 30% ethanol compared to 15% (“30% choice”). 49 animals were exposed to only water as controls for anxiety-related behavioral studies and RNAseq analyses, with consumption measured in mL on the treatment group’s IEA days. Daily drinking values were analyzed to identify outliers, leading to one daily value for one sample being removed.

**Figure 1.**
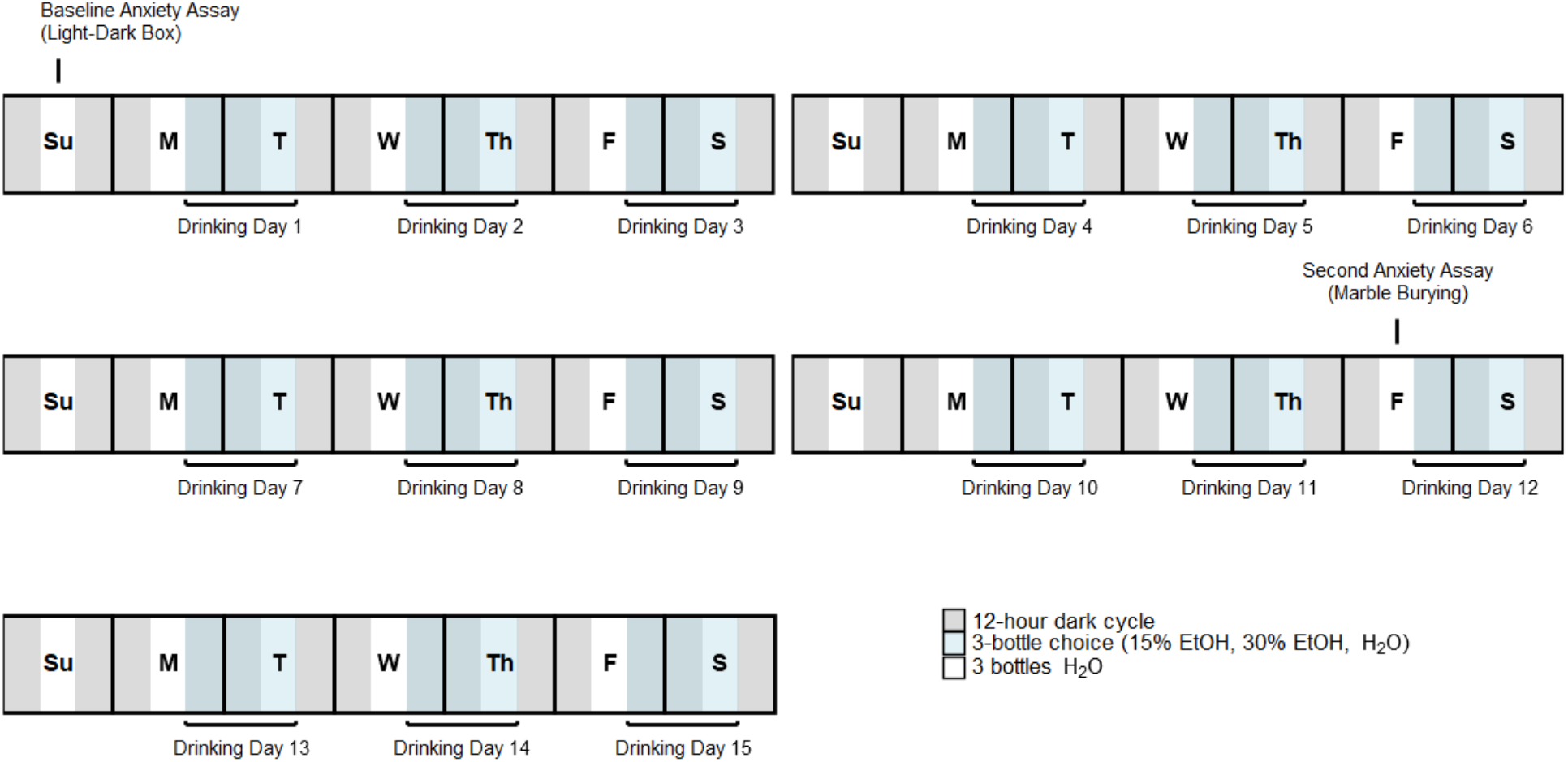
Timeline of intermittent ethanol access and anxiety-like behavioral assays. Mice were exposed to a three-bottle choice (H2O, 15% EtOH, 30% EtOH) drinking paradigm three days a week (Monday, Wednesday, and Friday) at the beginning of the dark cycle and fluid consumption measurements were taken after 24 hours.

Although not included in this report, all DO animals also underwent locomotor and anxiety-like behavioral testing via a light-dark box assay [26] 24 hours prior to initiation of IEA. Furthermore, after 4 weeks of IEA, animals were tested for anxiety-like behavior using a marble-burying assay at 24 hours after last access to ethanol. IEA consumption was then continued for 1 week (3 drinking days) prior to tissue harvesting as below. Only the first 4 uninterrupted weeks of IEA were used for the purposes of behavioral genetic studies in this report, excluding day 1 of IEA due to increased variance on initial exposure to ethanol bottles.

### Analysis of Behavioral Data

To identify the presence of any patterns in ethanol consumption, principal component analysis was conducted using daily ethanol consumption data in grams of ethanol per kilogram of body weight. Hierarchical clustering was carried out using Ward’s method on daily ethanol consumption data (in grams of ethanol per kilogram of body weight)[27]. Because of the presence of two superclusters which roughly divided drinking days into early (drinking days 2-7) and late (drinking days 8-13) phases, mean ethanol consumption, preference, and 30% choice phenotypes were derived for first week (drinking days 2-4), last week (drinking days 11-13), and whole study time periods (drinking days 2-13). Linear modeling of daily consumption and within-sample t-tests were used to compare first week and last week consumption and characterize progressive ethanol consumption across our mouse population.

### Tissue Sample Collection and Genotyping

Mice were sacrificed 24 hours after the end of their last ethanol exposure period via cervical dislocation and decaptitation and tissue samples were collected immediately afterwards, flash-frozen in liquid nitrogen and stored at -80° C. Tail snips were collected and sent to NeoGen Inc (Lincoln, NE) for DNA isolation and genotyping using a GigaMUGA microarray (*n*_SNP_ = 141,090; *n*_CNV_ = 2,169) designed to optimize genotyping of DO mice.[28] Mice were genotyped in two batches (*n*_1_ = 240; *n*_2_ = 420). Brains were microdissected into nine regions as previously described [29, 30]. Samples were only collected from 630 mice as 6 mice reached humane endpoints before the end of experimentation. After genotyping, 21 mice were removed due to low overall call quality (< 98% call rate), and one additional mouse was removed due to high heterozygosity (> 3 standard deviations above the mean) indicating contamination. Seven additional mice were removed due to potential sample mix-ups during data collection, resulting in a final sample of 603 mice (554 ethanol-drinking mice and 49 ethanol-naïve controls).

### Behavioral QTL Analysis

QTL mapping was carried out using the R/QTL2 software package for R. Dependent variables included whole study, first week, and last week averages for ethanol consumption, ethanol preference, and 30% choice [31]. SNPs were imputed based on progenitor haplotype probabilities at intervals of 0.01 Mbp and relatedness between individual mice was controlled for using a kinship matrix derived using the leave-one-chromosome-out (LOCO) method [32]. Cohort was included as a fixed-effect covariate in all of our analyses. Phenotypes were either log- (for consumption and 30% choice) or square-root-transformed (for ethanol preference) to obtain normality before running analyses. Genome-wide significance thresholds were derived using permutation analysis (*n*_perm_ = 1000), with an empirical threshold of *p* < 0.05 used to identify significant loci and a conventionally-used threshold of *p* < 0.63 used to identify suggestive loci. 95% Bayesian confidence intervals were identified for both significant and suggestive QTLs.

Positional candidate genes were identified as genes with coding regions located within significant bQTL confidence intervals using gene annotation databases from Mouse Genome Informatics (MGI).[33] To detect founder strain effects, haplotype analysis was conducted for chromosomes containing significant or suggestive bQTLs using best linear unbiased predictors (BLUPs) in R/QTL2. Lastly, LOD scores for individual SNP variants within significant bQTL confidence intervals were estimated using R/QTL2. The OrganismDbi package in R [34], together with information from MGI and ReMap [35], were then used to annotate known gene transcripts and regulatory elements from ChIP-seq, respectively, aligned to the GRCm38/mm10 genome assembly.

### RNA Extraction and Sequencing

Prefrontal cortex samples from 220 mice were chosen for RNA-seq analysis based on average total ethanol consumption during the fourth week of IEA. This included 100 mice from each extreme of the ethanol consumption distribution and an additional 20 ethanol-naïve control mice. Detailed analysis of RNA-seq data will be presented elsewhere. RNA was extracted and purified using RNeasy mRNA kits (Qiagen, Germantown, MD). Quality of isolated RNA was assessed using both a NanoDrop assay (Thermo Fisher Scientific, Waltham, MA) and BioAnalyzer 2100 (Agilent Technologies, Santa Clara, CA). Isolated RNA samples were then shipped on dry ice for poly-A selected library construction and sequencing at the Novogene Bioinformatics Institute (Sacramento, CA) as paired-end sequence 150 bp fragments with approximately 20 million reads per sample on the Illumina HiSeqX system.

RNA-seq quality control was performed using FastQC 0.11.7 and based on several metrics including base sequence quality, tile sequence quality, base sequence content, sequence length, duplication levels, and adapter content [36]. Trimmomatic 0.38 was used to trim the first 5 bp from the 3` end of each read and to trim the 5` end based on read quality [37]. After passing QC, a pooled transcriptome of all DO mouse founder strains was used to reconstruct and align to individual genomes for each subject using Genotype-by-RNA-Seq (GBRS)[38]. Finally, variant-stabilized transcript counts (VST) were quantified using DESeq2 [39].

### Expression QTL Analysis

QTL mapping was carried out using R/QTL2 software. VST-normalized GBRS expression data for individual gene transcripts were inserted as phenotypes, with the LOCO-derived kinship matrix and fixed-effect cohort covariates included in the analyses. Due to the large number of expression QTL (eQTL) analyses conducted, permutation analysis was not feasible for all eQTLs mapped; instead, a LOD threshold of 5 was used to remove likely insignificant eQTLs. eQTLs were further filtered to identify *cis-*eQTLs located < 2 Mbp from the correlated gene’s start or stop position and with 95% Bayesian confidence intervals < 2 Mbp in size. *D`* linkage disequilibrium estimates were then calculated between markers within confidence intervals of *cis-*eQTLs corresponding to positional candidate genes and peak bQTL markers [40].

### Bioinformatics Analysis of Candidate Genes

Variants including single nucleotide polymorphisms (SNPs), copy number variants/microsatellite regions, and insertions/deletions within significant bQTL confidence intervals were identified using the dbSNP annotation database from MGI [41]. dbSNP provides information on each variant’s predicted function class, defined by positional relatedness to a specific transcript for a gene. Variants within coding sequences and which change amino acid sequence, which are more likely to cause functional consequences to positional candidate genes, were identified using this database as well as variants with founder strain distributions matching observed haplotype patterns.

GWAS Catalog was searched for associations between human orthologs of positional candidate genes and alcohol-related traits [42]. Positional candidate genes were also used as search terms in GeneWeaver (http://geneweaver.org) to identify relevant data sets related to ethanol or other substance use [43].

## Results

### DO Mice Exhibit Progressive Increase in Ethanol Consumption Over Time

The population of 600 DO mice exhibited wide variation in ethanol consumption, preference, and 30% choice over 5 weeks of IEA (Figure S1). First and last week consumption were significantly correlated (0.58, *p* = 1.08×10^−52^), suggesting that highly stable factors may contribute to observed variance in consumption across the study (Figure 2). Preference was significantly positively correlated with ethanol consumption across all time periods (*p* < 0.05). 30% choice was not significantly correlated with ethanol consumption across all time periods, suggesting that mice which prefer a higher concentration of ethanol do not consume significantly different amounts of ethanol overall.

**Figure 2.**
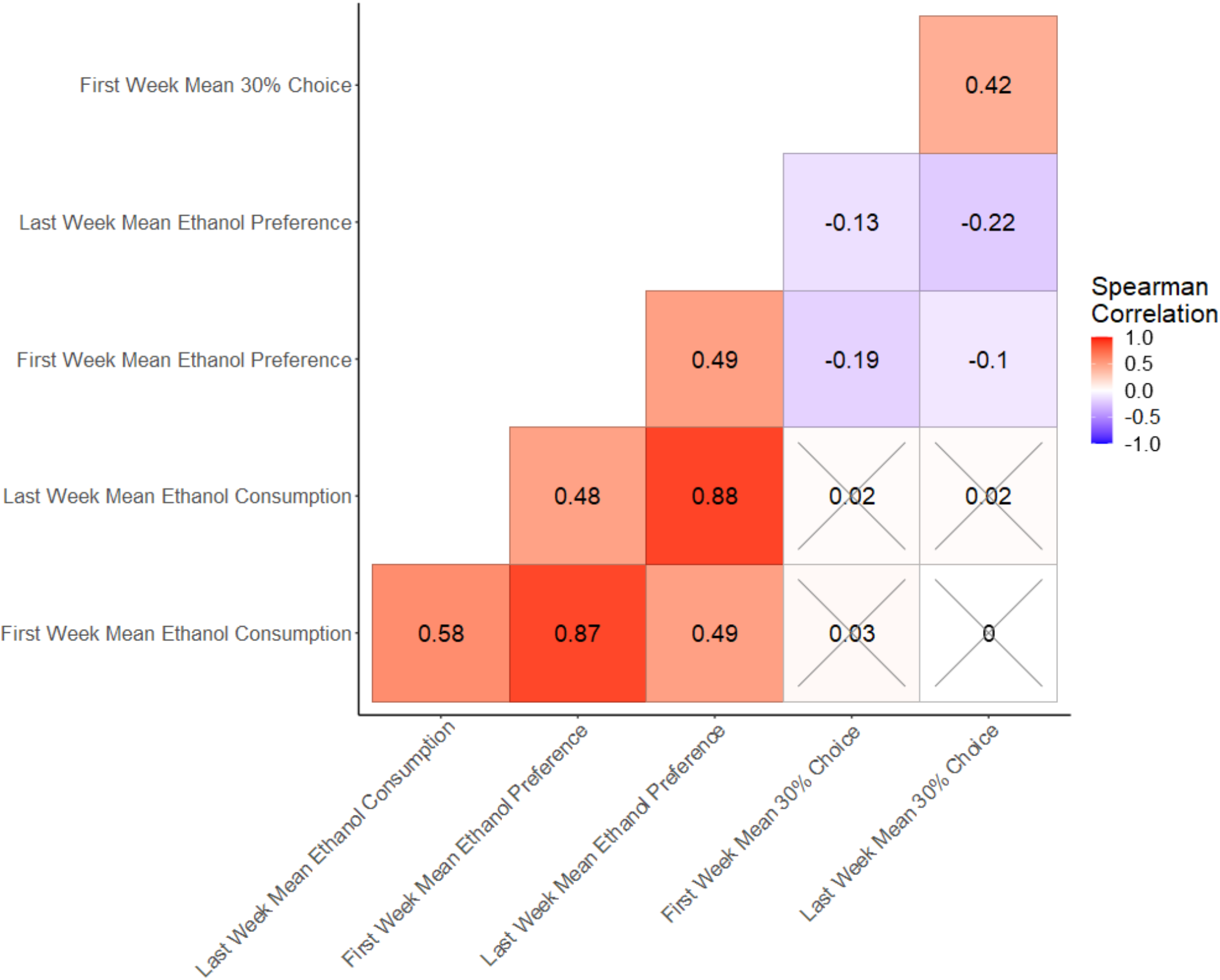
Spearman correlations between ethanol consumption behaviors. Ethanol consumption, preference, and 30% choice were measured over 4 weeks of IEA via three-bottle choice. Spearman correlations were calculated for first week and last week phenotypes. First and last week means were significantly positively correlated within each phenotype; preference also significantly positively correlated with consumption for each time period. 30% choice was negatively correlated with preference, but not significantly correlated with overall consumption. Crosses denote insignificant correlations (*p* > 0.05).

To determine potential substructure within the ethanol consumption time course, we performed hierarchical clustering of daily ethanol consumption. This generated two superclusters, one containing drinking days 2-7 and one containing drinking days 8-13 (Figure 3A). Despite the strong correlation noted above, ethanol consumption in the last week of the study was significantly higher than in the first week (Figure 3B, *p* = 1.006×10^−05^), as confirmed by the progressive increase in ethanol consumption over the course of the study (Figure 3C).

**Figure 3.**
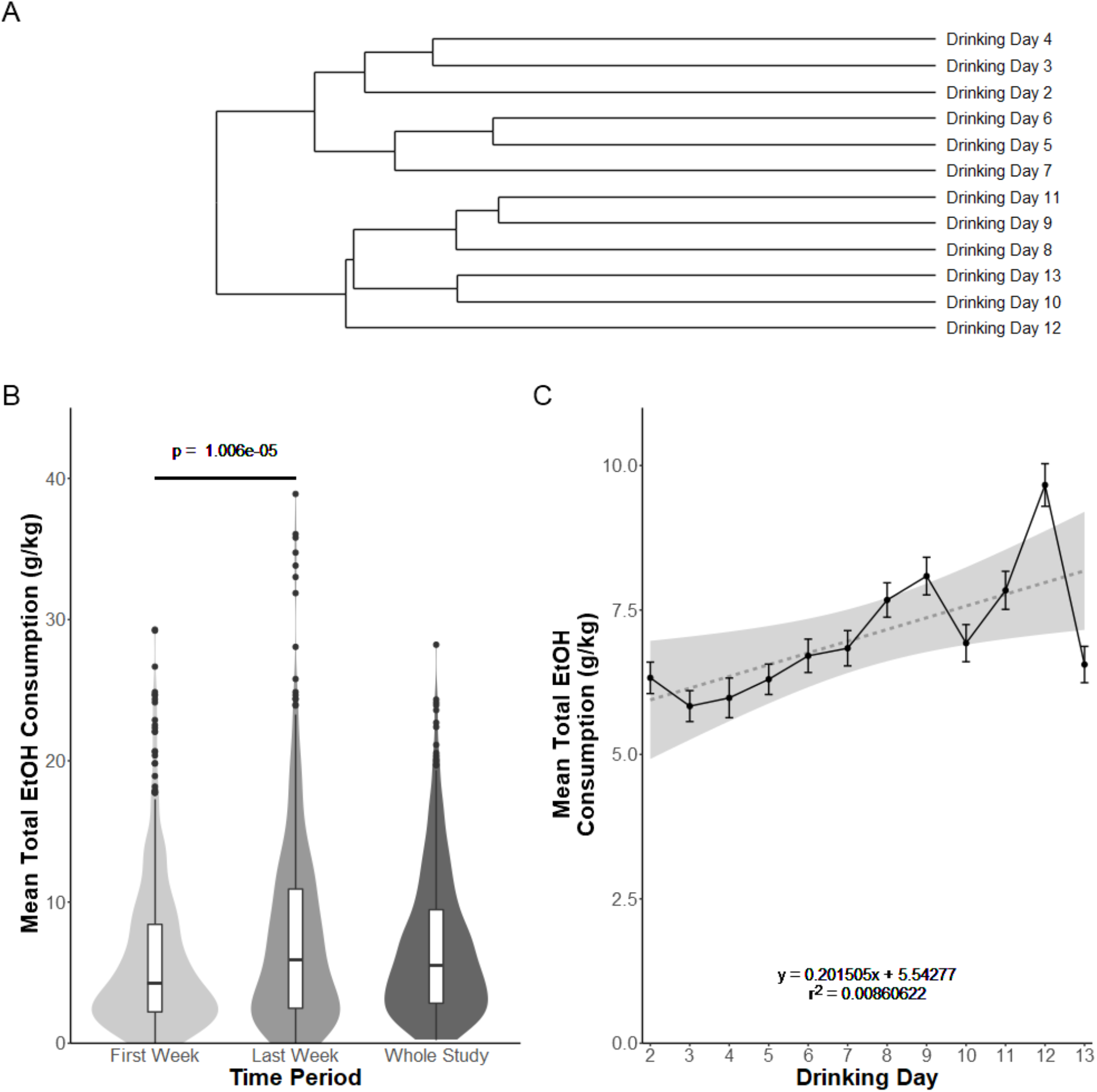
Diversity Outbred mice demonstrate progressive increase in ethanol consumption over five weeks of intermittent ethanol access. Hierarchical clustering of daily ethanol consumption identified two superclusters, one including drinking days 2-7 and one including drinking days 8-13 (A). Last week mean ethanol consumption was significantly higher than first week mean consumption in a within-sample one-tailed t-test (B). Drinking day was a significant predictor of ethanol consumption in linear regression, with consumption increasing by approximately 0.2 g EtOH/kg body weight each successive day (C).

Principal component analysis identified that one principal component likely explained a large percentage (42.4%) of the variance in daily ethanol consumption (Figure S2A) The first principal component appeared highly correlated with whole study mean total ethanol consumption, (Figure S2B) while a second principal component correlated positively with first week mean ethanol consumption (Figure S2C) and negatively with last week mean ethanol consumption (Figure S2D). While a single principal component may explain overall ethanol consumption, the second component appears to explain variance in first and last week consumption suggesting that there may be additional genetic factors controlling the amount of escalation in consumption over time.

SNP-based heritability estimates for ethanol consumption ranged from 0.197-0.310, and estimates for ethanol preference ranged from 0.143-0.301. Generally, first week heritability estimates were lower than either last week or whole-study estimates. Estimates for 30% choice were much lower, ranging from 0.00-0.073 (Table 1).

**Table 1.**
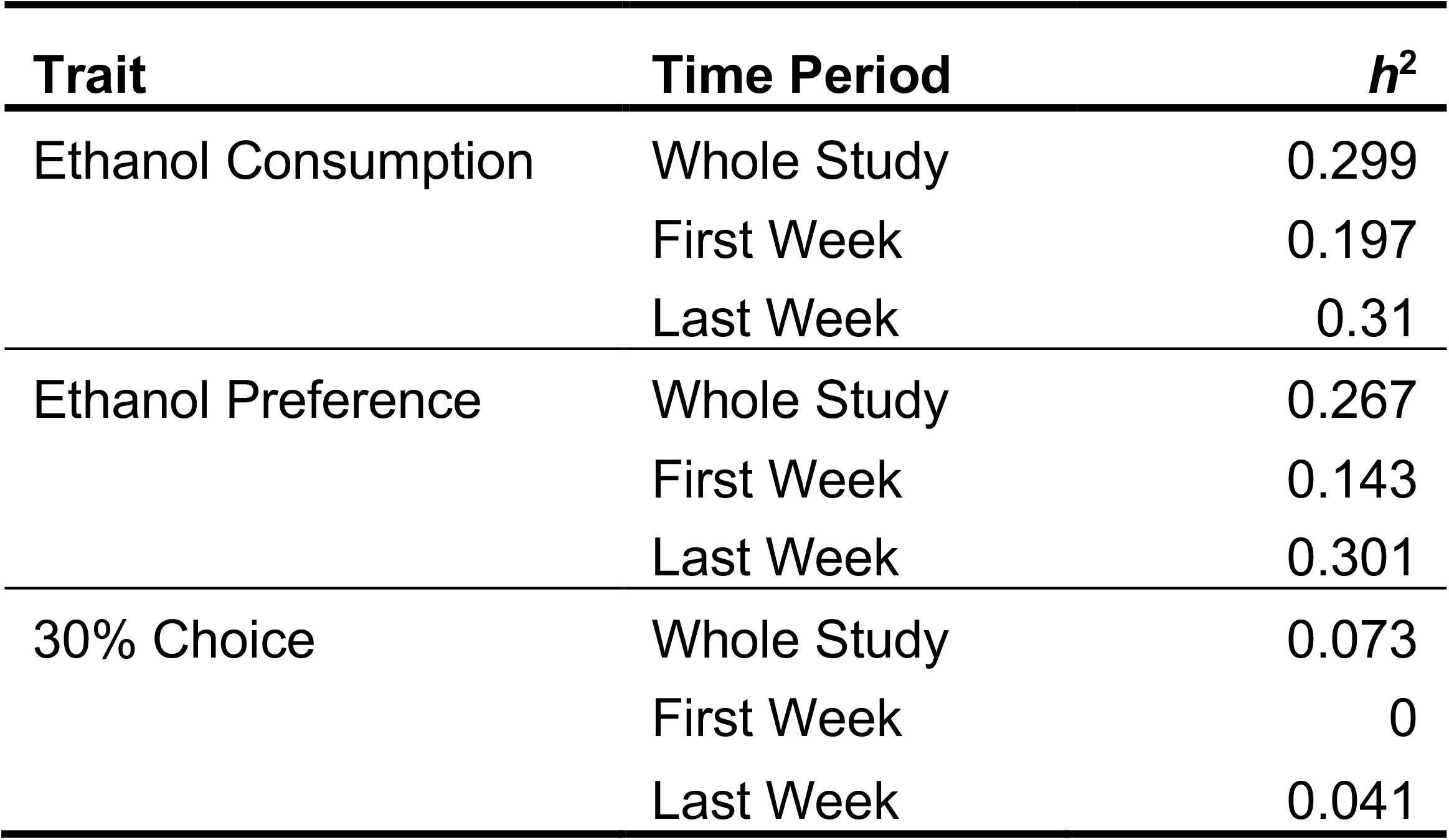
Ethanol consumption phenotypes are heritable in our DO mouse population. Heritability estimates were estimated using restricted maximum likelihood in R/QTL2. Consumption and preference phenotypes both had heritability estimates ranging from 0.14-0.31; 30% choice had lower heritability estimates (< 0.08).

| **Trait** | **Time Period** | **$h^2$** |
| --- | --- | --- |
| Ethanol Consumption | Whole Study | 0.299 |
|  | First Week | 0.197 |
|  | Last Week | 0.31 |
| Ethanol Preference | Whole Study | 0.267 |
|  | First Week | 0.143 |
|  | Last Week | 0.301 |
| 30% Choice | Whole Study | 0.073 |
|  | First Week | 0 |
|  | Last Week | 0.041 |

### bQTL Analysis Identifies 3 Significant Loci

Significant bQTLs (*p* < 0.05) were identified on Chromosome (Chr) 4 which explained 5.79% of variance in last week mean ethanol consumption (LOD = 8.23), on Chr 3 which explained 6.06% of variance in last week mean 30% choice (LOD = 8.63), and on Chr 12 which explained 5.3% of variance in first week mean ethanol preference (LOD = 7.52) (Figure 4). An additional 12 suggestive QTLs (*p* < 0.63) were identified, and Bayesian confidence intervals were estimated for all QTLs (Table 2). All suggestive bQTLs explained between 4-5% of observed variance in their relevant phenotype. A total of 1,375 genes were located within significant or suggestive confidence intervals and thus designated as provisional positional candidate genes.

**Figure 4.**
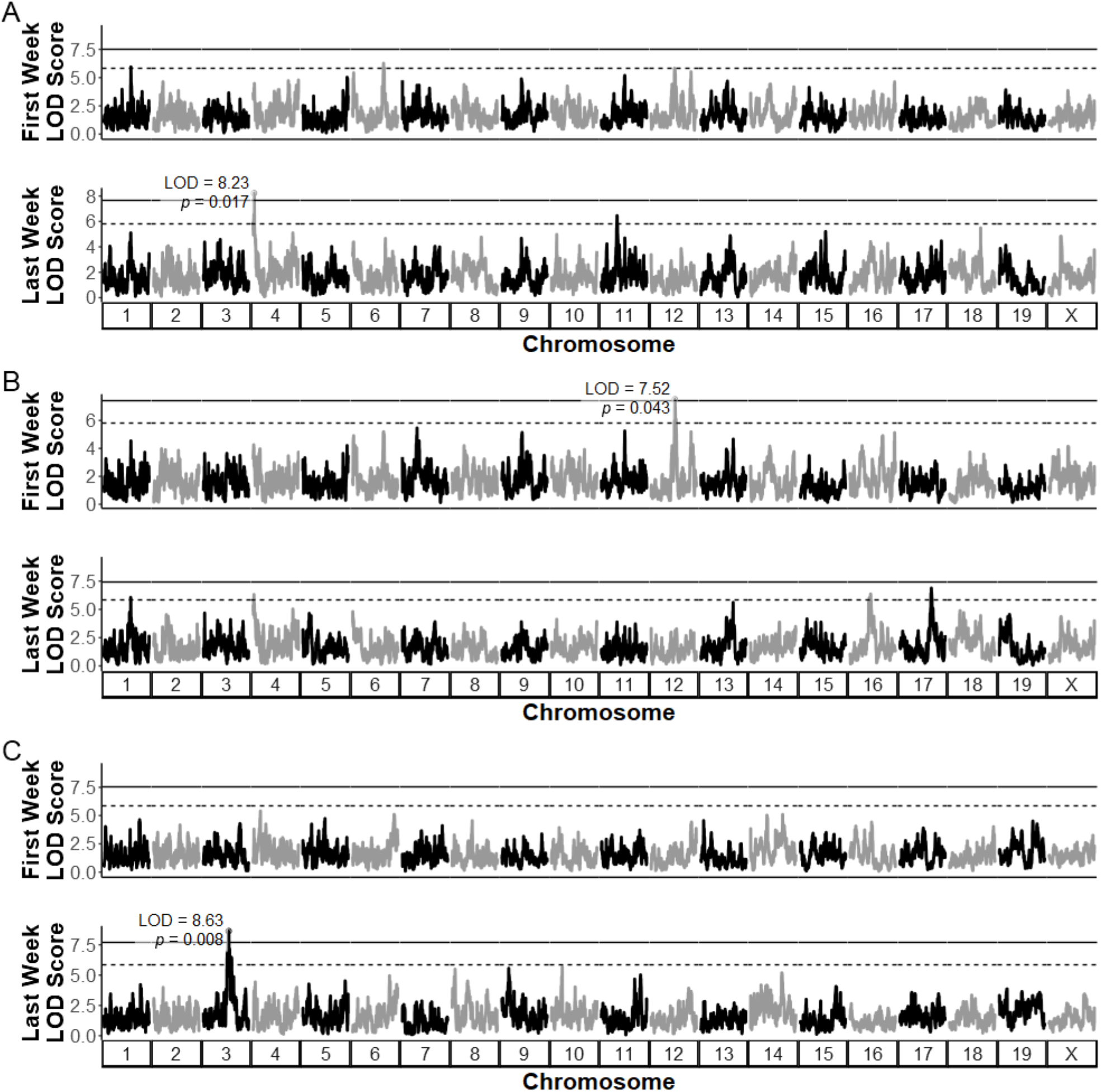
QTL analysis identifies significant peaks for three ethanol-related behavioral phenotypes. Different genetic effects were identified for first week (top) and last week (bottom) ethanol consumption (A), preference (B), and 30% choice (C) behavioral QTLs. For ethanol consumption, no significant QTL was observed during the first week of the study, but a significant QTL on Chromosome 4 was identified in the last week (LOD = 8.23; *p* = 0.017). For ethanol preference, a significant QTL was observed on Chromosome 12 (LOD = 7.52, *p* = 0.043) during the first week, but not the last week of the study. For 30% choice a significant QTL was identified on Chromosome 3 during the last week (LOD = 8.63, *p* = 0.008), but not the first week of the study. Empirical significance thresholds for each phenotype were calculated using permutation analysis (n_perm_ = 1000); solid black lines represent *p* < 0.05 and dashed black lines represent *p* < 0.63 thresholds.

**Table 2.** Suggestive and significant QTLs for ethanol consumption behaviors. A population of 636 Diversitty Outbred (DO) mice were exposed to five weeks of intermittent ethanol access via a 3-bottle choice paradigm. QTLs were identified for first week, last week, and whole study mean ethanol consumption, preference for ethanol compared to water (“ethanol preference”), and preference for 30% ethanol compared to 15% ethanol (“30% choice”). QTLs are annotated with peak LOD scores, positions in megabase pairs (Mbp), 95% Bayesian confidence intervals (C.I.), and the number of genes identified using MGI gene annotations within each C.I.

| Phenotype | LOD Score | p-value | Chr | Peak Marker Position (Mbp) | 95% Bayesian C.I. (Mbp) | C.I. Length (Mbp) | Genes | % Variance Explained |
| --- | --- | --- | --- | --- | --- | --- | --- | --- |
| Last Week Mean 30% Choice | 8.63 | 0.008 | 3 | 108.23 | 105.92-109.69 | 3.77 | 149 | 7.13 |
| Last Week Mean Ethanol Consumption | 8.23 | 0.017 | 4 | 8.41 | 8.24-9.29 | 1.05 | 20 | 6.8 |
| First Week Mean Ethanol Preference | 7.52 | 0.043 | 12 | 79.35 | 78.09-79.91 | 1.82 | 44 | 6.25 |
| Last Week Mean 30% Choice | 6.93 | 0.146 | 3 | 103.57 | 101.85-104.47 | 2.62 | 96 | 5.77 |
| Last Week Mean Ethanol Preference | 6.88 | 0.16 | 17 | 74.32 | 72.91-75.21 | 2.3 | 55 | 5.72 |
| Whole Study Mean Ethanol Consumption | 6.67 | 0.217 | 4 | 8.32 | 3.78-9.65 | 5.88 | 84 | 5.56 |
| Whole Study Mean Ethanol Preference | 6.65 | 0.227 | 1 | 135.35 | 134.64-136.59 | 1.94 | 70 | 5.54 |
| Last Week Mean 30% Choice | 6.48 | 0.278 | 3 | 118.93 | 118.39-119.81 | 1.42 | 19 | 5.41 |
| Last Week Mean Ethanol Consumption | 6.46 | 0.294 | 11 | 50.58 | 49.96-51.52 | 1.56 | 62 | 5.38 |
| Last Week Mean Ethanol Preference | 6.36 | 0.335 | 15 | 44.69 | 39.43-45.68 | 6.25 | 89 | 5.3 |
| Last Week Mean Ethanol Preference | 6.32 | 0.351 | 4 | 8.32 | 3.37-11.05 | 7.68 | 123 | 5.27 |
| Whole Study Mean Ethanol Preference | 6.3 | 0.362 | 4 | 7.00 | 3.37-8.32 | 4.95 | 73 | 5.26 |
| Whole Study Mean Ethanol Consumption | 6.29 | 0.363 | 11 | 70.61 | 69.86-71.34 | 1.48 | 104 | 5.25 |
| First Week Mean Ethanol Consumption | 6.27 | 0.378 | 6 | 113.92 | 112.74-115.31 | 2.56 | 90 | 5.23 |
| Last Week Mean Ethanol Preference | 6.06 | 0.47 | 1 | 134.73 | 119.53-144.91 | 25.38 | 544 | 5.05 |

Haplotype analysis within Rqtl2 was performed to implicate patterns of strain-specific allelic contributions to variation in ethanol traits at the significant bQTLs. For the significant locus with last week mean ethanol consumption on Chr 4, A/J alleles were associated with lower last week mean ethanol consumption, whereas NOD and PWK alleles correlated with increased last week ethanol consumption (Figure 5A). For the locus on Chr 3, A/J, C57BL/6J, and WSB/EiJ alleles correlated with an increase in 30% choice whereas PWK/PhJ alleles associated with a decrease in this phenotype (Figure 5B). NZO/HlLtJ and A/J alleles at the significant locus on Chr 12 were associated with an increase in first week ethanol preference, while CAST/EiJ, NOD/SHiLtJ, and WSB/EiJ alleles correlated with decreased ethanol preference (Figure 5C).

**Figure 5.**
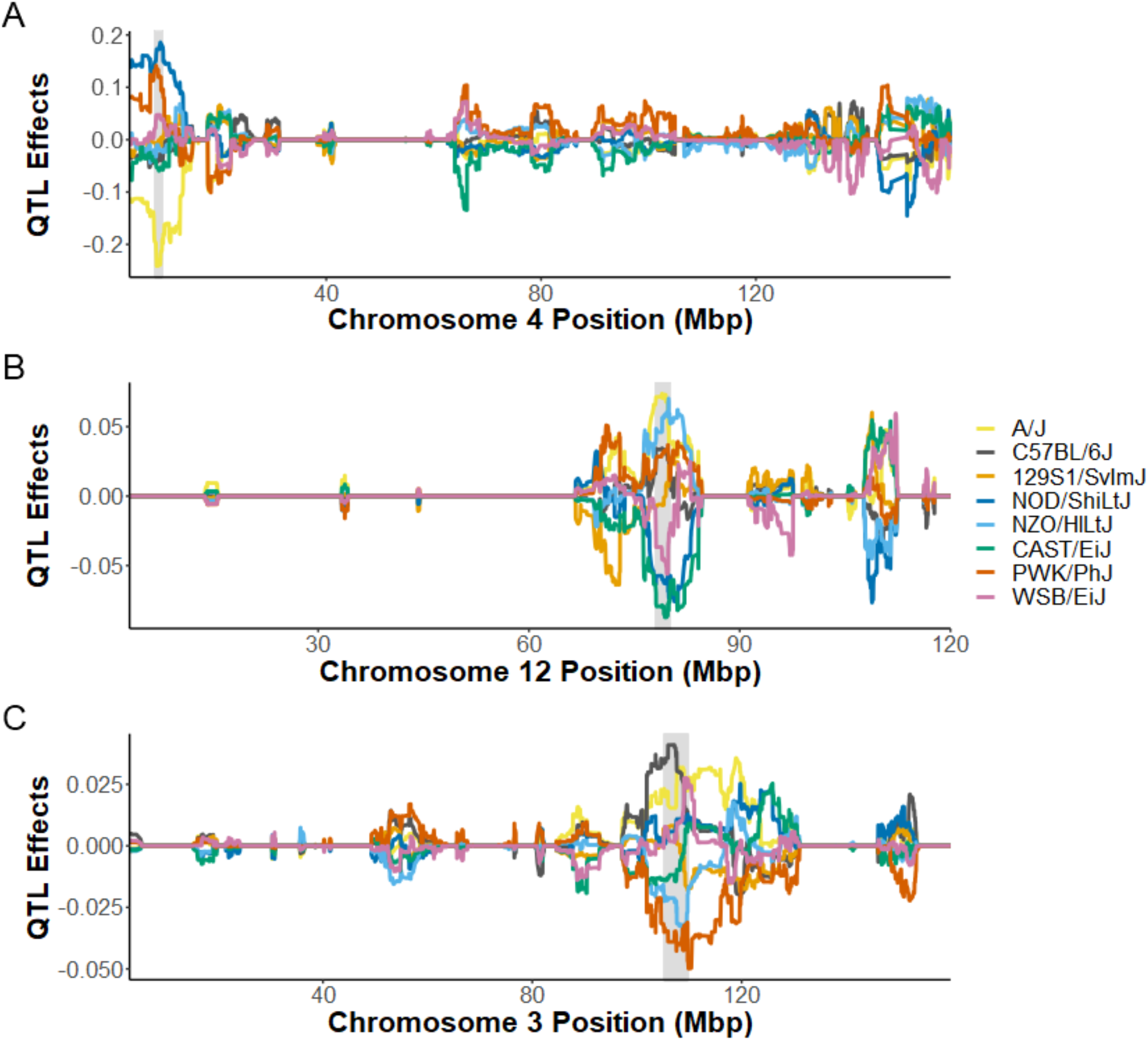
Haplotype analysis of significant QTLs identifies effects of DO progenitor alleles on ethanol consumption phenotypes. Best linear unbiased predictors (BLUP) were used to reduce signal noise and estimate effects of alleles from each of the 8 DO mouse founder strains; 95% Bayesian confidence interval around significant bQTL is indicated by the shaded background. A/J alleles (yellow) within the Chromosome 4 CI correlated with a reduction in last week ethanol consumption, whereas NOD (dark blue) and PWK (red) alleles correlated with observed increases in last week ethanol consumption (A). A/J alleles also correlated with an increase in ethanol preference for the CI on Chromosome 12, while NOD and CAST (green) alleles correlated with a decrease in ethanol preference for that locus (B). For the chromosome 3 locus, C57BL/6J (dark gray) and A/J alleles correlated with an increase in 30% choice, whereas PWK and NZO (light blue) alleles correlated with a decrease in 30% choice (C).

### Top variants identified within significant bQTL intervals

Top variants were identified as those within 1.5 LOD units from the highest variant LOD score in an interval (Figure 6A). For the significant bQTL on Chr 4, the variant with the highest LOD score was rs249655952 (LOD = 3.84), a downstream variant for predicted gene *Gm37386*. All other top variants for this region (*n* = 7) were intergenic variants (Table 3). This suggests that the variants which are causal for the bQTL on Chr 4 may be more involved in the regulation of gene expression than in causing functional changes to protein structures encoded by genes in the region; in fact, despite the presence of relatively few known gene transcript annotations for this region (Figure 6B), a number of regulatory elements have been identified and annotated across this region using ChIP-seq databases (Figure 6C). Among these top variants, 5 were unique to the A/J progenitor strain. The remaining 2 were intergenic deletions, both around 100 bp, unique to NOD/ShiLtJ. Haplotype analysis of this locus (Figure 5A) also suggests a role for alleles from these two strains, identifying that A/J alleles across the bQTL interval are correlated with lower last week ethanol consumption and that NOD/ShiLTJ alleles correlated with higher last week ethanol consumption.

**Figure 6.**
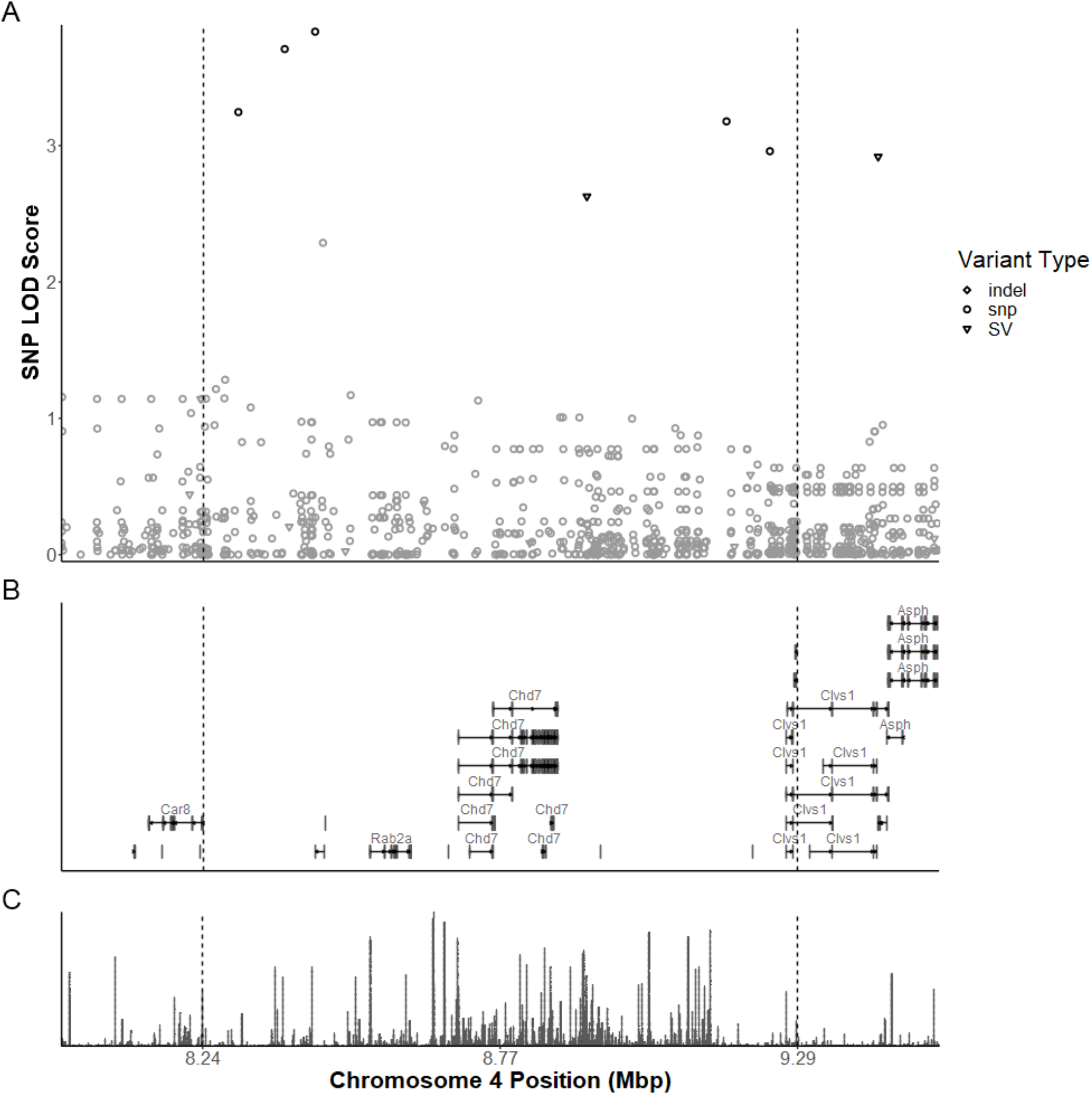
Variant LOD scores across significant bQTL on chromosome 4 for last week mean ethanol consumption. SNP associations were estimated across the 95% Bayesian confidence interval identified for the significant bQTL, identified by dashed vertical lines (A). Top variants (black) were identified as those with a LOD score within 1.5 of the highest score are black; all others are gray. Few known gene transcript annotations exist within this C.I. (B), but include *Rab2a, Chd7*, and *Clvs1*. Despite the relatively small number of genes in this interval, ReMap regulatory element density reveals a number of regulatory elements across the interval (C).

**Table 3.** Top SNPs from significant bQTL for last week mean ethanol consumption on Chromosome 4. LOD scores were calculated for individual variants across the 95% Baytsian C.I. for the significant bQTL. Top variants were identified as those within 1.5 LOD units of the top-scoring variant. Of these, none are coding sequence variants for identified genes, suggesting an important role of transcriptional regulation for this QTL.

| Variant | Type | Position (Mbp) | Alleles (Reference\|Variant) | Consequence | SNP LOD | Strain with Variant Allele |
| --- | --- | --- | --- | --- | --- | --- |
| 4:8302313_A/C | SNP | 8.302313 | A\|C | Intergenic variant | 3.25 | A/J |
| 4:8384190_G/A | SNP | 8.38419 | G\|A | Intergenic variant | 3.71 | A/J |
| rs249655952 | SNP | 8.437965 | T\|A | Gm37386 downstream variant | 3.84 | A/J |
| SV_4_8917963_8918080 | Structural Variant | 8.918022 | -\|4:8917963-8918080;DEL;RAW;DEL | NA | 2.63 | NOD/ShiLtJ |
| rs32455519 | SNP | 9.164587 | C\|T | Intergenic variant | 3.18 | A/J |
| rs262840752 | SNP | 9.241249 | C\|T | Intergenic variant | 2.96 | A/J |
| SV_4_9432463_9432570 | Structural Variant | 9.432516 | -\|4:9432463-9432570;DEL;RAW;DEL | NA | 2.92 | NOD/ShiLtJ |

Similarly, variant LOD scores, transcripts, and regulatory elements from both the Chr 12 C.I. for first week ethanol preference (Figure S3) and the Chr 3 C.I. for last week 30% choice (Figure S4) were identified. Within the Chr 12 interval, 23 variants were identified within 1.5 LOD units of the top-scoring variant (rs47944281, LOD = 6.00). Intron variants made up 11 of these, while 7 were intergenic variants, 3 were downstream variants, 1 was an upstream variant, and 1 (rs51613251, LOD = 4.52) was a missense variant in *Zfyve26*. This variant alters amino acid sequence from in alanine the reference sequence (i.e., C57BL/6J and all other progenitor strains) to glutamine in CAST/EiJ and NOD/ShiLtJ mice. This is consistent with the findings of the haplotype analysis (Figure 5B), in which both strains decreased first week mean preference for ethanol; however, WSB/EiJ alleles also conferred decreased first week mean preference.

For the Chr 3 interval, 250 variants were identified within 1.5 LOD units of the top scoring variant (rs49087152, LOD = 6.13). Again, the majority of variants had no functional consequence for amino acid sequence; however, 1 variant (rs29605696, LOD = 5.10) was a missense variant for *Celsr2* and another variant (rs253441023, LOD = 4.69) introduced a stop codon in *Sypl2*. In the NZO/HILtJ, CAST/EiJ, and PWK/PhJ progenitor strains, the missense variant in *Celsr2* alters amino acid sequence from leucine in the reference sequence (i.e., C57BL/6J and all other progenitor strains) to proline [33]. Our haplotype analysis is consistent with the strain distribution pattern of this variant, demonstrating a decrease in last week 30% choice corresponding to alleles from all three progenitor strains in which the proline amino acid is present (Figure 5C). The premature stop codon in *Sypl2* is present in NZO/HILtJ, CAST/EiJ, PWK/PhJ, and WSB/EiJ founder strains; however, WSB/EiJ alleles correspond with an increase in last week 30% choice, making the haplotype pattern inconsistent with the founder strain distribution for the rs253441023 variant.

### eQTL Analysis

As an initial step to aid ranking positional candidate genes, eQTL analysis on RNAseq data from prefrontal cortex was performed and analyzed using GBRS to allow optimal alignment and allele-specific expression estimation [38], with the assumption that cis-eQTLs could indicate genetic variance in gene expression contributing to the identified bQTL. This analysis identified 186,840 markers with eQTLs having LOD scores > 5. Of these, 9,107 loci were identified as *cis*-eQTLs for 7,635 genes. We further filtered these results to identify *cis*-eQTLs overlapping bQTL confidence intervals (Table 4).

**Table 2.**
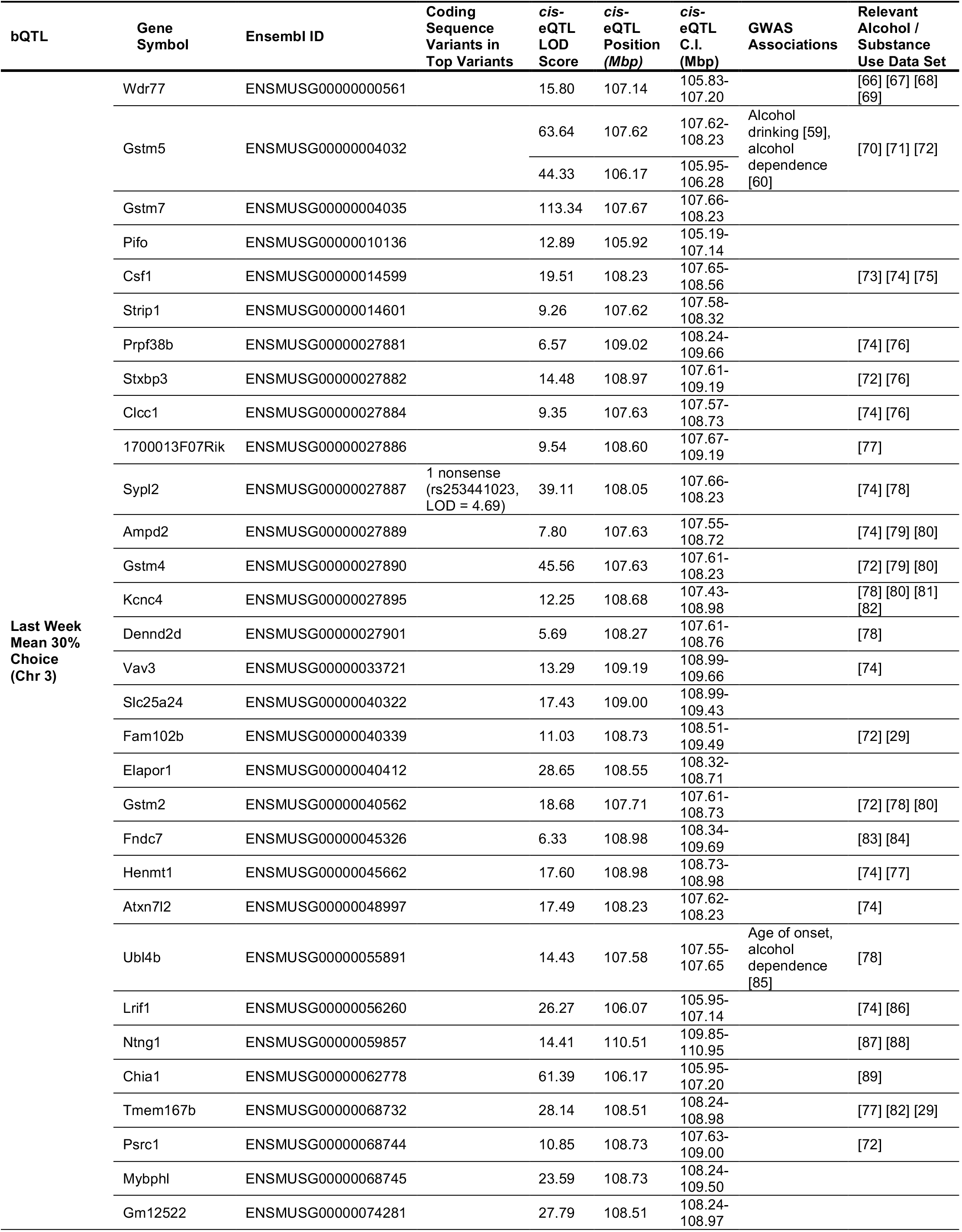
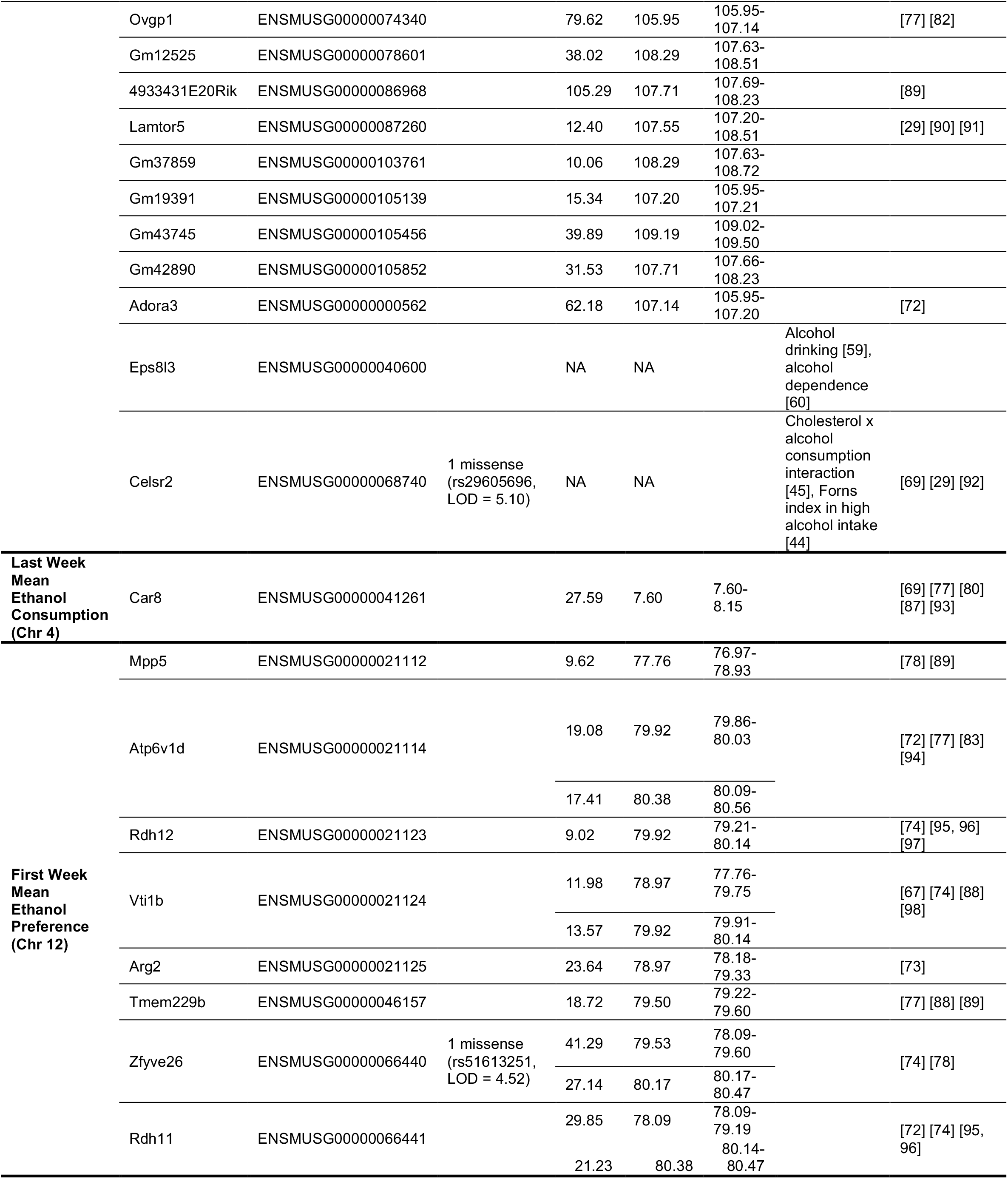
Top candidate genes from significant bQTL confidence intervals. Positional candidate genes were identified as having coding regions within significant bQTL intervals using the MGI gene annotation database. Top candidate genes were identified as those with either a cis-eQTL from prefrontal cortex RNA-seq (n = 220) overlapping the relevant bQTL C.I. or with a significant human GWAS association from GWAS Catalog. Top genes were then used as search queries on GeneWeaver to identify presence within relevant alcohol or substance use data sets.

Notably, only one gene within the significant bQTL confidence interval for last week mean ethanol consumption on Chr 4 had a *cis*-eQTL which met filtering criteria: *Carbonic anhydrase 8* (*Car8*) (Figure 7A). Haplotype analysis of the *Car8 cis-*eQTL indicated that A/J alleles at the locus correlate with increased expression of *Car8* in prefrontal cortex; this suggests that variants within this locus on Chr 4 may confer not only an increase in *Car8* expression, but a decrease in ethanol consumption during the last week of the study (Figure 7B). *Car8* expression had a significant negative Spearman correlation with both last week ethanol consumption (*r* = -0.22; *p* = 0.008) and last week ethanol preference (*r* = -0.23; *p* = 0.006), consistent with the relationship suggested by the effect of A/J alleles. Taken together, these results suggest a role for variants unique to A/J mice in both regulation of *Car8* expression and ethanol consumption in mice.

**Figure 7.**
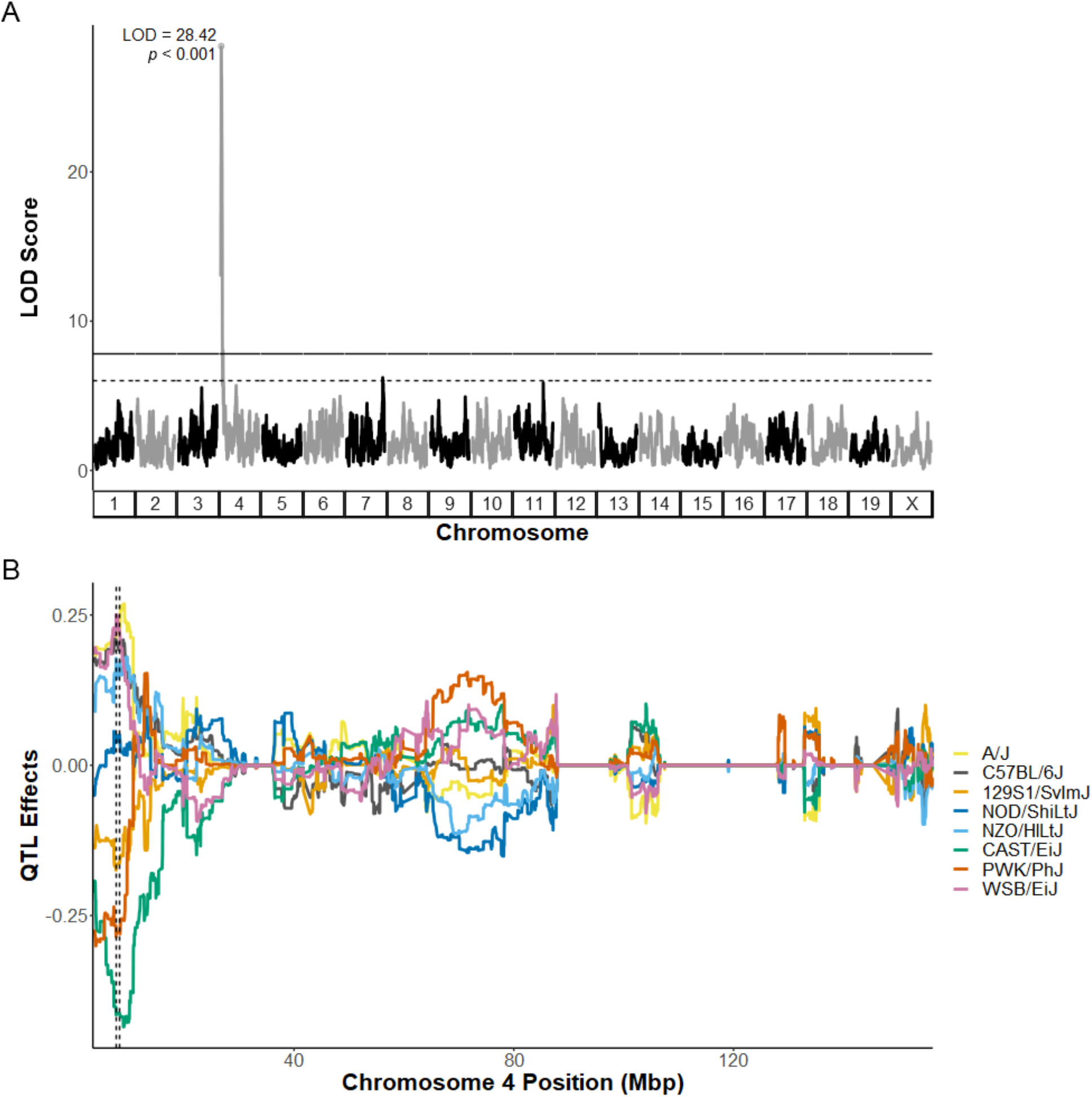
*Car8* eQTL overlaps significant bQTL for last week mean ethanol consumption on chromosome 4 and displays effects of A/J alleles. Expression QTL analysis using prefrontal cortex samples identifies *Car8* as the only gene with a significant *cis*-eQTL (LOD = 28.42; *p* < 0.001) overlapping the significant bQTL for last week ethanol consumption (A). Haplotype analysis reveals that mice with A/J alleles (yellow) at this locus display increased expression of *Car8* (B).

The significant bQTL for last week mean 30% choice had 40 genes with cis-eQTLs with LOD scores > 5 (Table 4). Of these, only *Sypl2* contained coding sequence variation, while 2 genes also had significant alcohol-related GWAS associations from GWAS Catalog (*Gstm5* and *Ubl4b*). Notably, *Gstm5* had two corresponding cis-eQTL peaks, located approximately 1.5 megabases apart. No *cis*-eQTL was observed for *Celsr2* at this locus, meaning that this variation in coding sequence—and not regulation of the gene’s expression—would be a likely mechanism for this candidate gene’s role in ethanol-related phenotypes. Twelve genes within the significant bQTL C.I. for first week mean ethanol preference, including *Zfyve26*, had cis-eQTLs with LOD scores > 5. Only *Zfyve26* had a coding sequence variant in the top variants, and none of the genes at this locus had significant alcohol-related GWAS associations from GWAS Catalog.

### Bioinformatics Analysis Identifies 6 Genes with Significant Human GWAS Associations

Only 4 genes from significant bQTLs were identified as having significant associations with alcohol-related traits in the GWAS Catalog database. (Table 4) Of these, one gene was related to glutathione metabolism (*Gstm5*), while two appeared related to epidermal growth factor pathways (*Eps8l3* and *Celsr2*) and the final gene is part of a transmembrane family (*Tm4sf5*). An additional 2 genes located within a suggestive bQTL for first week mean ethanol consumption on Chr 6 also had significant associations from GWAS Catalog, *Slc6a1* and *Slc6a11*, both members of a GABA transporter family.

In GWAS Catalog, *Celsr2* was associated with Forns index (a measure of liver fibrosis) in individuals with high alcohol intake [44] and with interaction terms between alcohol consumption and both LDL and HDL cholesterol levels [45]. As mentioned above, this gene also has a missense variant within the DO mouse population that ranked as one of the top variants within the significant QTL for last week 30% choice on Chr 3.

## Discussion

This report is the first to analyze genetic variance in ethanol consumption using DO mice. The design of these studies provided an opportunity to analyze a progressive ethanol consumption phenotype at high genetic resolution in a model assessing a large proportion of the genetic variance across mouse populations. Our results show progressive ethanol consumption across the population but a suggestion of differing genetic influences on initial ethanol intake (first week) versus chronic consumption (last week). We identified 3 significant and 12 suggestive bQTLs spread across a total of 7 chromosomes. In most cases, the support intervals for these bQTL in DO mice were much narrower (Table 2, median = 2.3 Mbp) than seen in any prior mouse genetic study on ethanol consumption. The integration of cis-eQTL and bioinformatic analyses allowed prioritization of an experimentally tractable number of high priority candidate genes, most of which have not been previously implicated in mechanisms or risk factors underlying ethanol consumption.

### bQTL Analysis Identifies Specific, Novel Loci for Ethanol Consumption Behaviors

Our high-resolution genetic mapping and candidate gene identification for ethanol consumption, ethanol preference, and 30% ethanol choice, identified 3 significant bQTLs (Figure 4): one on Chr 4 associated with last week ethanol consumption, one on Chr 3 associated with last week 30% choice, and one on Chr 12 associated with first week ethanol preference. A total of 12 suggestive bQTLs were also identified on Chrs 1, 3, 4, 5, 11, 15, and 16 (Table 2); notably, the suggestive bQTLs on Chr 4 for whole study ethanol consumption, last week ethanol preference, and whole study ethanol preference directly overlap the significant QTL for last week ethanol consumption but have broader Bayesian confidence intervals. This may suggest that variants within that locus influence not only ethanol-drinking behaviors in the last week of IEA, but also across the study. This would not be unexpected given the significant correlations between last week and whole-study measures, as with both ethanol consumption and preference phenotypes.

Some of these bQTLs overlapped regions implicated in prior in mouse genetic model analysis of ethanol behaviors. For example, all 3 bQTLs on Chr 3 are located within the confidence interval for alcohol preference identified by Belknap and Atkins in 2001 [11, 46]. However, our QTL confidence intervals are an order of magnitude smaller, with the broadest being the significant interval for last week 30% choice at approximately 3.77 Mbp, compared to 62.18 Mbp for the Belknap and Atkins QTL. The genes in this region were found in a dataset from a rat ethanol consumption QTL [47] and a rat alcohol sensitivity QTL [48], suggesting an overlap with a syntenic locus involved in ethanol-related phenotypes in rat models. Similarly, our suggestive QTL for whole study ethanol consumption on Chr 11 is a similar, but more specific locus as an earlier QTL identified by Kirstein et al. in 2002 [49]. Here our confidence interval was only 1.48 Mbp compared to 20.95 Mbp. These results demonstrate that the use of the DO mouse allows for identification of novel loci modulating ethanol consumption at a much higher resolution that was previously possible.

### eQTL Analysis Identifies Car8 as a Top Candidate Gene

We used the presence of a *cis-*eQTL as a step in annotating our candidate gene lists as the presence of an eQTL in the proximity of a bQTL (and with markers in linkage disequilibrium with the peak bQTL marker) provide strong evidence that genetic variation in a region is modifying both observed differences in ethanol-related behaviors and expression of a candidate gene. One example of such a gene is *Car8*, located on the proximal end of the significant bQTL for last week ethanol consumption on Chr 4 (Figure 6). *Car8* has a *cis-*eQTL with markers in strong linkage disequilibrium with the peak marker of the bQTL; furthermore, haplotype analysis reveals that A/J alleles at this locus correlate both with decreased ethanol consumption in the last week of IEA and increased expression of *Car8* in the prefrontal cortex. Additionally, the lack of top SNPs in the Chr 4 confidence interval with functional consequences affecting protein structure suggest that the variants within that region responsible for the observed bQTL are involved in the regulation of gene expression. We consider these data strong Sort for *Car8* as a priority candidate gene for ethanol consumption.

*Car8* shares sequence similarity to the carbonic anhydrase family of genes, but lacks carbonic anhydrase activity; instead, it is known to bind to and inhibit the IP3R1 calcium signaling channel [50]. IP3R1 has been suggested to play a role in ethanol-enhanced GABA release in cerebellum, a potential mechanism by which ethanol-induced cerebellar ataxia occurs [51]. *Car8* is highly expressed in cerebellar Purkinje cells and implicated in both cerebellar development and locomotor function [52, 53]. However, *Car8* is also expressed significantly across multiple other brain regions (see: http://mouse.brain-map.org/experiment/show/191). *Car8* has also been show to be regulated by ethanol within cerebellum, ventral midbrain and anterior cingulate in mice (Table 4). Given this ethanol regulation in brain, the significant negative correlations between *Car8* expression and last week ethanol consumption, and the strong *Car8* eQTL in linkage dysequilibrium with the Chr 4 bQTL for last week ethanol consumption, we hypothesize that altered expression of *Car8* may modulate ethanol consumption.

The significant bQTL for last week mean 30% choice on Chr 3 had four notable genes, including *Celsr2* and *Sypl2*, which both had coding sequence variants, and *Eps8l3* and *Gstm5*, which had association data in GWAS Catalog with alcohol-associated traits (Table 4). *Celsr2* encodes a transmembrane protein and has been implicated in axon development in the forebrain[54] and regulation of motor neuron regeneration following injury [55]. *Sypl2* encodes a synaptophysin-like protein, but remains relatively uncharacterized; however, it has been implicated in obesity [56, 57] and major depressive disorder [58] in human GWAS literature. Both of these genes would require further study to characterize any effects on alcohol-related behaviors. *Gstm5* and *Eps8l3* had GWAS associations with alcohol-associated traits in GWAS catalog through a small number of SNPs mapped to both transcripts, rs6693815, rs11102001, and rs11102002 [42], including an interaction between fibrinogen levels and alcohol consumption [59] and alcohol dependence measures [60], although neither gene reached genome-wide statistical significance for either study.

Within the significant bQTL for first weak ethanol preference on Chr 12, only *Zfye26* had both a cis-eQTL with a LOD score > 5 and a coding sequence variant, a missense variant. *Zfye26* has been implicated in hereditary spastic paraplegia [61-63] through a role in autophagy [64, 65], but little is known about a potential role in alcohol consumption, dependence, or other ethanol-related behaviors in animal models.

Due to the large number of identified intergenic variants within significant bQTL intervals, we plan to further analyze expression data collected from our prefrontal cortex samples to collect additional support for candidate genes, as well as to characterize individual strong candidates more thoroughly. Further studies on *Car8* are underway due to the evidence supporting it as a strong candidate.

## Conclusions

Genetic mapping using the DO mouse led to identification of three significant bQTLs for ethanol consumption behaviors with narrow confidence intervals (1-4 Mbp). This high-resolution genetic mapping allowed for identification of viable candidate genes for ethanol consumption behaviors. Furthermore, incorporation of RNA-seq data from prefrontal cortex samples collected from the same population of mice and integration with human GWAS data and cross-species data from GeneWeaver generated robust annotation and prioritization of positional candidate genes. These findings will characterize genes and mechanisms causing observed differences in ethanol consumption in mice, helping lead to a better mechanistic understanding of the complex genetic architecture of ethanol consumption and alcohol use disorder in humans.

## Acknowledgements

The authors thank members of the Miles Laboratory for their support during the performance of these analyses. Nicholas Rodriguez was instrumental in performing GBRS analyses. We also thank Drs.

Vivek Philip and K.B. Choi at The Jackson Laboratory for assistance in utilizing GBRS for RNAseq analysis.

## Author Contributions

Conceived and designed the study: KM LM MFM. Acquired the data: KM LM. Analyzed the data: ZT KM ZS JN MM. Wrote the paper: ZT KM ZS MS MFM.

---

**Supplemental Figure 1.**
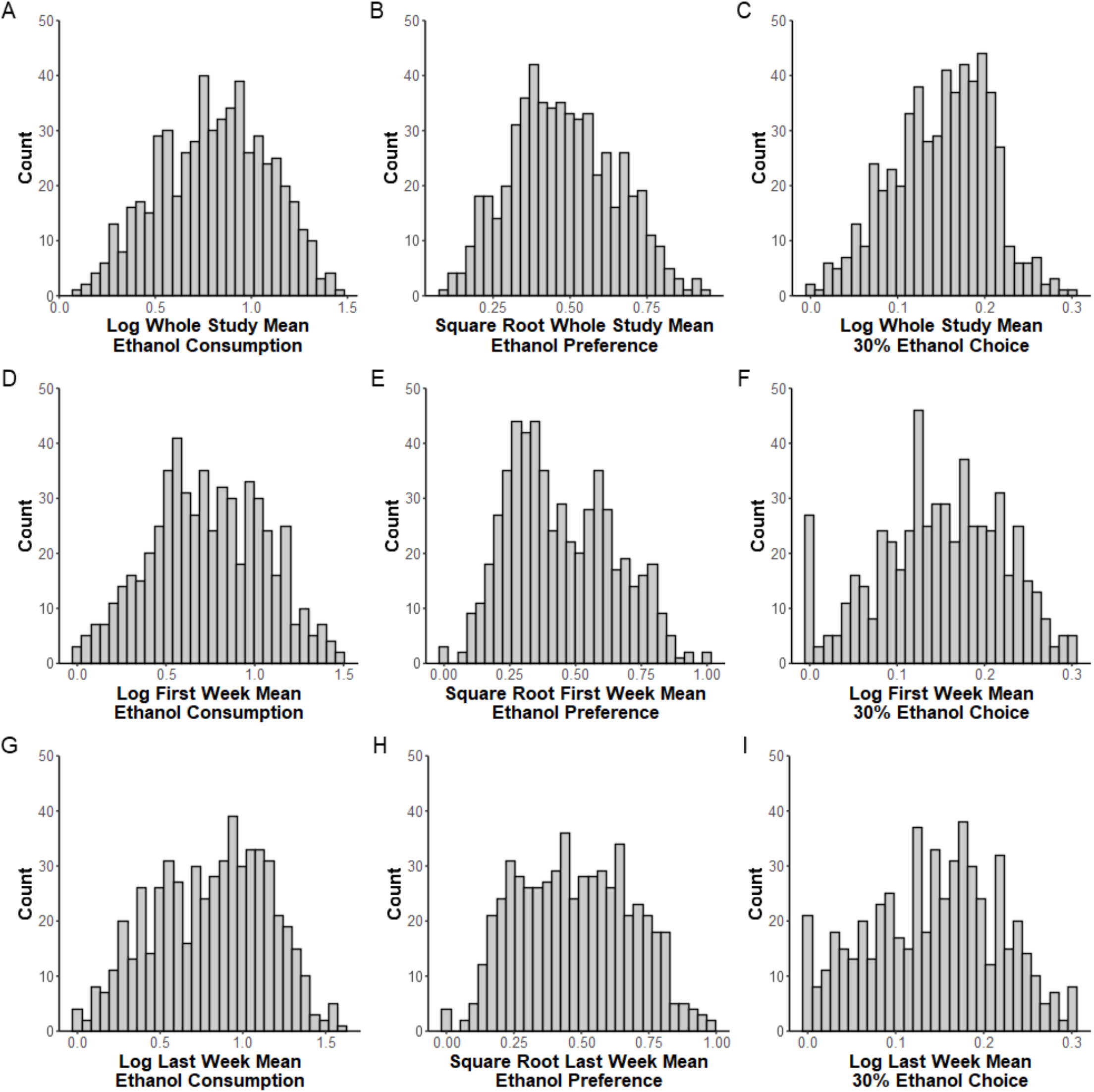
Histograms of whole study, first week, and last week ethanol consumption, preference, and 30% choice following normalization show variability in ethanol drinking phenotypes across DO mice. First week (drinking days 2-4), last week (drinking days 11-13), and whole study (drinking days 2-13) mean consumption values were calculated for each mouse (*n* = 636). Consumption values were then used to calculate preference for ethanol over water (“ethanol preference) and preference for a higher concentration of ethanol compared to total ethanol (“30% ethanol choice”). Consumption and 30% ethanol choice were log-transformed for normalization; preference was square root-transformed.

**Supplemental Figure 2.**
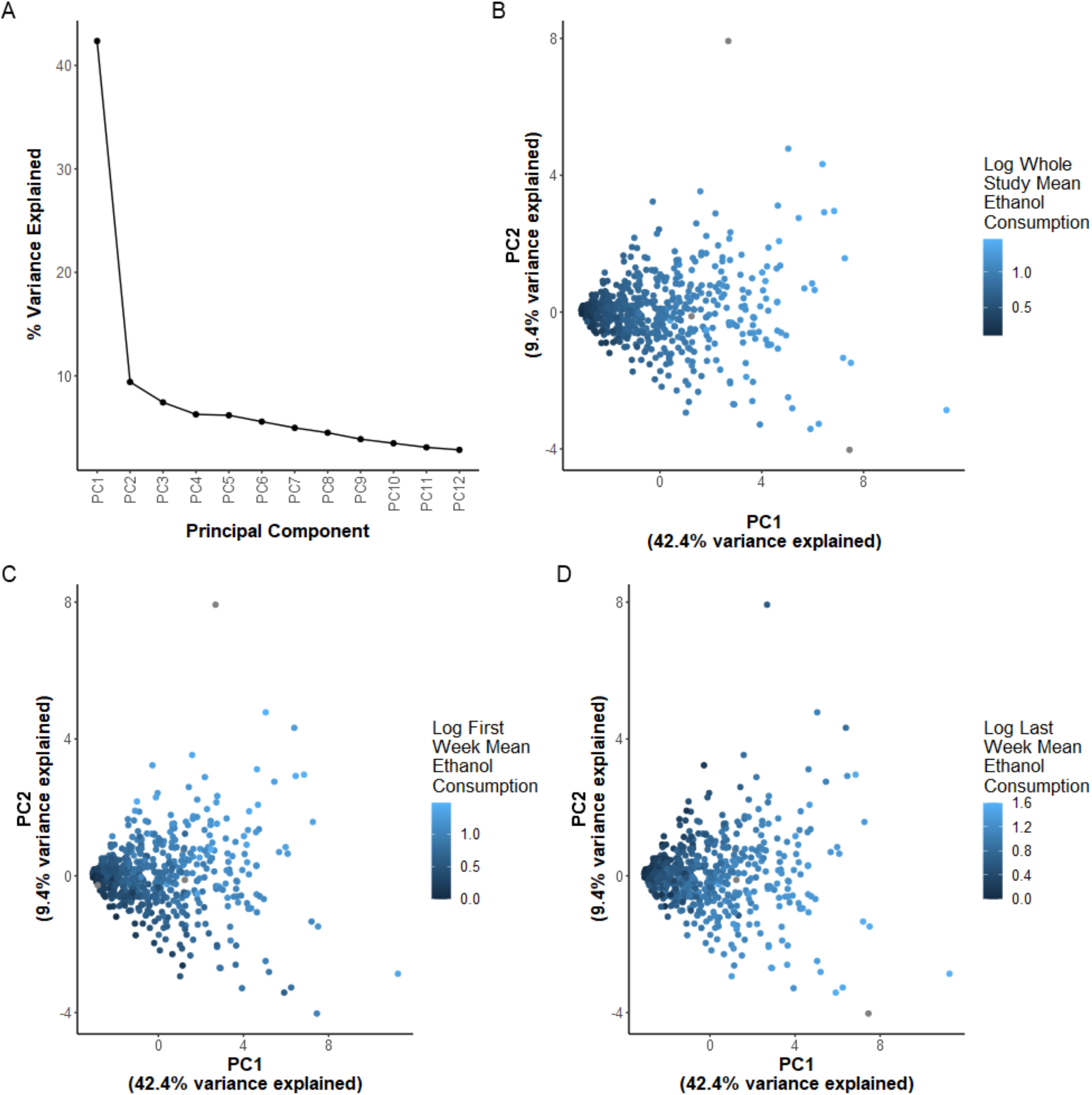
Principal component analysis of daily drinking values across DO mice reveals main effect of whole study mean ethanol consumption. A single principal component explained 42.4% of the total variance in daily drinking across 636 male DO mice (A). This component appears correlated with whole study mean ethanol consumption (B), while principal component 2 appears positively correlated with first week mean ethanol consumption (C) and negatively correlated with last week mean ethanol consumption (D).

**Supplemental Figure 3.**
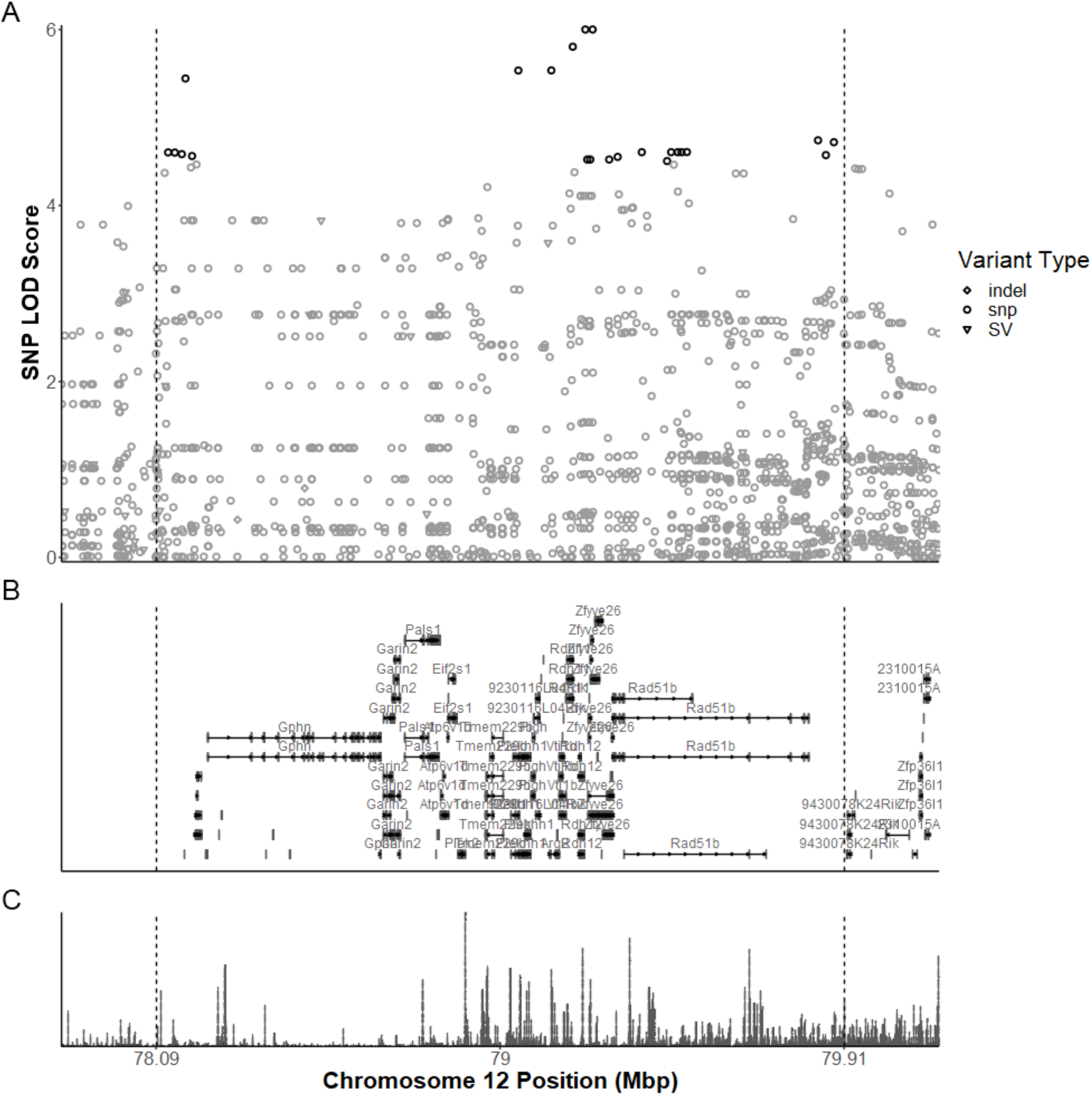
Variant LOD scores across significant bQTL on chromosome 12 for first week mean ethanol preference. SNP associations were estimated across the 95% Bayesian confidence interval identified for the significant bQTL, identified by dashed vertical lines (A). Top variants (black) were identified as those with a LOD score within 1.5 of the highest score are black; all others are gray. Known gene transcript annotations exist within this C.I. include *Zfyve26* (B). ReMap regulatory element density reveals a number of regulatory elements across the interval (C).

**Supplemental Figure 3.**
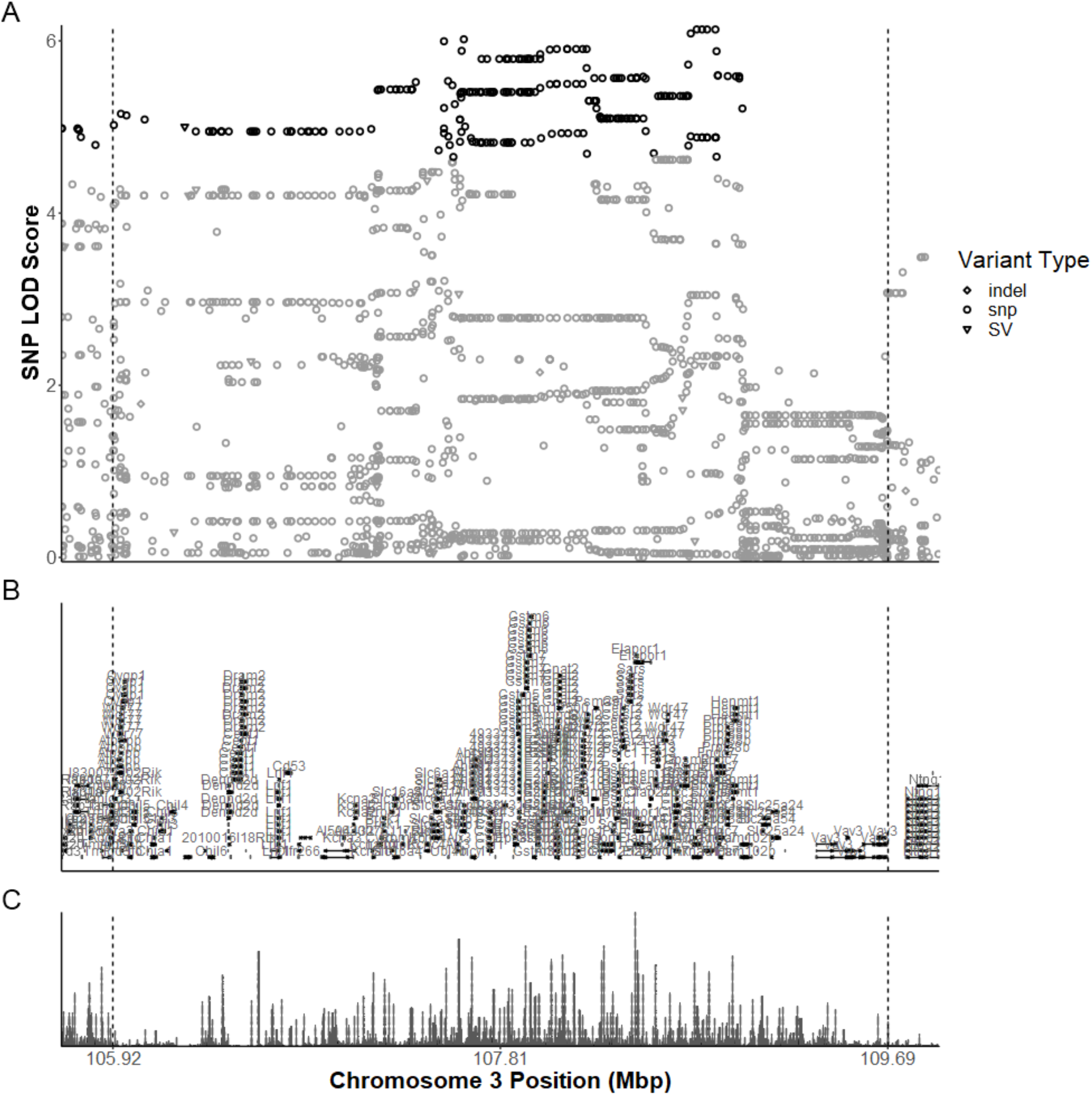
Variant LOD scores across significant bQTL on chromosome 3 for last week mean 30% ethanol choice. SNP associations were estimated across the 95% Bayesian confidence interval identified for the significant bQTL, identified by dashed vertical lines (A). Top variants (black) were identified as those with a LOD score within 1.5 of the highest score are black; all others are gray. Known gene transcript annotations exist within this C.I. include *Celsr2* and *Sypl2* (B). ReMap regulatory element density reveals a number of regulatory elements across the interval (C).

